# Identification of oxalyl-CoA synthetase gene (*LsAAE3*) and its regulatory role in β-ODAP biosynthesis in grasspea (*Lathyrus sativus* L.)

**DOI:** 10.1101/2021.04.23.441068

**Authors:** Neetu Singh Kushwah, P.S. Shanmugavadivel, Alok Das, Meenal Rathore, Archana Singh, Narendra Pratap Singh

**Affiliations:** ICAR-Indian Institute of Pulses Research, Kanpur-208 024, India

**Keywords:** β-ODAP, CoA-dependent pathway of oxalate degradation, Expression analysis, Grasspea/*Lathyrus sativus*, *LsAAE3*, Oxalyl-CoA synthetase

## Abstract

Grasspea is a popular pulse crop due to its hardiness and low cost of production. Presence of anti-nutritive factor ‘β-ODAP’ in its seeds and other plant parts hinder its widespread cultivation and usage. Oxalyl-CoA synthetase is one of the key enzyme of β-ODAP biosynthesis pathway, catalyses the conversion of oxalate to oxalyl-CoA. *ACYL ACTIVATING ENZYME 3* (*AAE3*) gene has been characterised to encode an oxalyl-CoA synthetase enzyme in many plant species. We report here the isolation of full length *AAE3* homolog in grasspea with a combination of PCR based strategy and *in silico* analysis. We first identified *AAE3* homolog by PCR using degenerate primers. The partial Ls*AAE3* sequence showed 88% amino acids sequence identity with the characterised *AAE3* gene of *M. truncatula*. We then predicted the full length *AAE3* sequence using the publically available transcriptome datasets of grasspea. Determination of *LsAAE3* gene and protein structure and phylogenetic relationship analysis strongly suggested that *LsAAE3* is a true homolog of *AAE3* gene. Expression profiling of *LsAAE3* in grasspea varieties with contrast in β-ODAP content revealed its inverse relationship with the β-ODAP content, *LsAAE3* thus negatively regulates the synthesis of β-ODAP. Involvement of AAE3 encoded oxalyl-CoA synthetase in a CoA-dependent pathway of oxalate degradation is well proven in many plant species. We also identified the CoA-dependent pathway of oxalate degradation in grasspea. Based on these observations, we hypothesized that *LsAAE3* may regulate β-ODAP content, possibly, by CoA-dependent pathway of oxalate degradation in grasspea. If this hypothesis is substantiated, genetic manipulation of *LsAAE3* presents viable option for reducing β-ODAP content in grass pea.

## Introduction

*Lathyrus sativus* (Family: Fabaceae) commonly known as grasspea is a grain legume crop, cultivated for food, feed and fodder purposes mainly in South Asia and East Africa in an area of about 1.5 million hectares with an annual production of 1.20 million tons (Lambein et al. 2019). Due to its remarkable ability to tolerate drought as well as excess precipitation, it is mainly cultivated in drought-prone and rainfed areas where extreme weather conditions are prevalent. Together with, its capability to grow on a wide variety of soils, including very poor to heavy clay soils, and the ability to fix atmospheric nitrogen makes it an attractive crop for adverse climatic conditions (Campbell 1997). Despite having these climate-smart features, its cultivation is limited due to the presence of anti-nutritional factor β-ODAP in its seeds and other plant parts which causes a neurodegenerative disease, neurolathyrism, on continuous consumption of grasspea seeds in a large quantity. Complete removal of β-ODAP or reduction in its amount will aid in realizing the full potential of this crop in the face of climate change. Absence of β-ODAP-free lines in the grasspea germplasm or wild *Lathyrus* spp. (Campbell et al. 1987; Abd-El-Moneim et al. 2001; Kumar et al. 2011) and instability of low-ODAP trait under extreme weather conditions (Fikre et al. 2008) are the major bottlenecks in the development of stable low/ β-ODAP-free grasspea varieties by conventional breeding. This has necessitated the need of exploring candidate genes encoding the enzymes of β-ODAP biosynthesis and its regulation and eventually utilizing them for developing β-ODAP-free grasspea varieties using modern biotechnological tools. Published data suggests that the ODAP biosynthesis pathway involves oxalyl-CoA synthetase and ODAP synthase enzymes (Malathi et al. 1970). Oxalyl-CoA synthetase catalyses the conversion of oxalate to oxalyl-CoA, which then acts as one of the substrates for the ODAP synthase enzyme to catalyse the formation of β-ODAP (Fig. 1a) (Malathi et al. 1970).

**Fig. 1.**
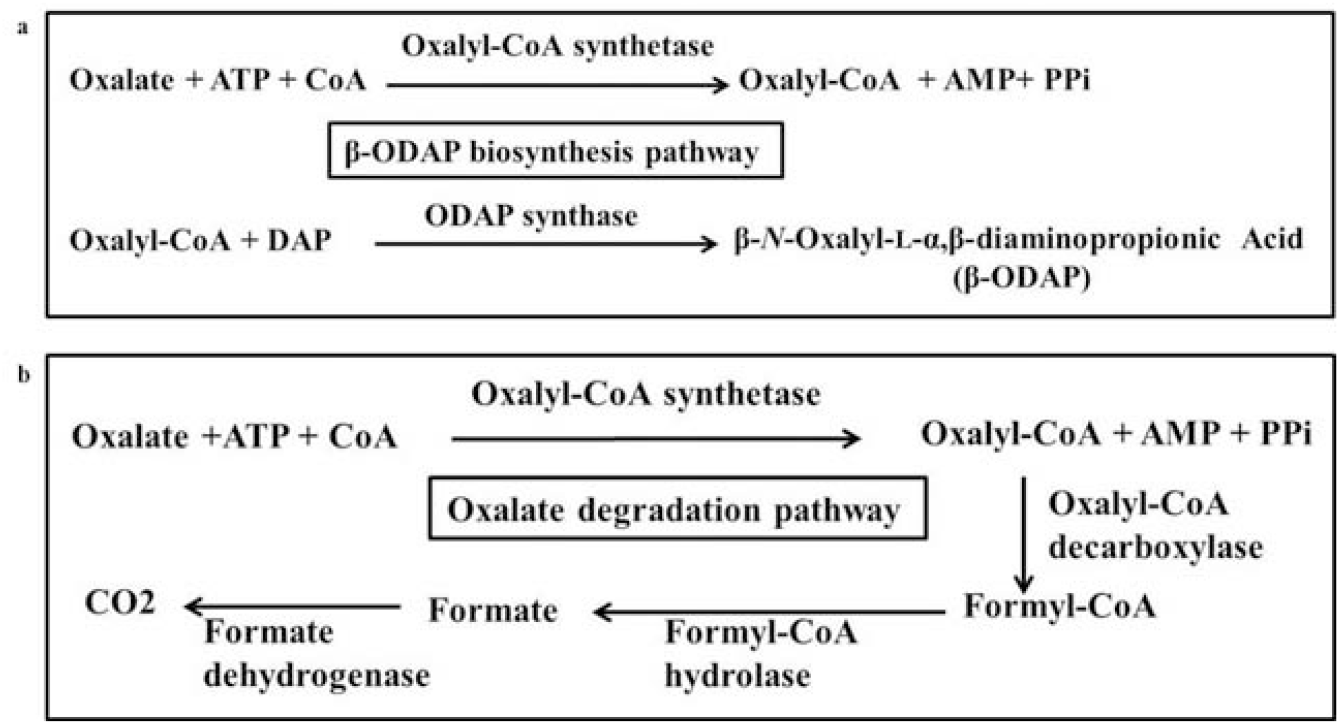
Schematic representation of proposed ODAP biosynthesis and CoA-dependent pathway of oxalate catabolism in plants. **a** ODAP biosynthesis pathway in grasspea. **b** CoA-dependent oxalate degradation pathway in plants

Biochemical activity of oxalyl-CoA synthetase has been reported in grasspea (Malathi et al. 1970) as well as in several other legume crops (Adsule and Barat 1977). However, no information of genes encoding oxalyl-CoA synthetase enzyme were available until 2012. The *Acyl Activating Enzyme 3* (*AAE3*) gene was first time discovered to function as an oxalyl-CoA synthetase in *Arabidopsis thaliana* (Foster et al. 2012). Later on, the *AAE3* ortholog was reported in the budding yeast *Saccharomyces cerevisiae* (Foster and Nakata 2014), model legume *Medicago truncatula* (Foster et al. 2016), rice bean (Lou et al. 2016) and rice (Peng et al. 2017). In all these non-ODAP forming plant species, oxalyl-CoA synthetase enzyme was reported to be involved in proposed multi-step oxalate degradation pathway (Foster et al., 2012; Giovanelli and Tobin; Fig. 1b) where oxalate is converted to oxalyl-CoA by Oxalyl-CoA synthetase, which is then decarboxylated to form formyl-CoA followed by formate and eventually into CO2 by the activity of a group of enzymes including Oxalyl-CoA decarboxylase, Formyl-CoA hydrolase, and Formate dehydrogenase. The genes encoding other enzymes of the CoA-dependent pathway of oxalate degradation have also been identified in *Arabidopsis* and maize (Foster et al. 2012; Yang et al. 2018). However, in grasspea, neither oxalyl-CoA synthetase gene nor the other components of CoA-dependent pathway of oxalate degradation has been reported so far. Here, we report the identification of the candidate gene encoding oxalyl-CoA synthetase based on homologous gene sequence information, its characterization and expression pattern in different plant parts, and different varieties of grasspea varying in β-ODAP content.

## Materials and methods

### Plant materials and growth conditions

The low-ODAP (cv. Mahateora) and medium-ODAP (cv. Pusa-24) containing varieties of grasspea were procured from ICAR-IIPR, Regional Station, Bhopal (MP), India. Seeds were sown in pots filled with autoclaved vermiculite to grow for 15 to 20 days in the culture room, maintained at 23±2°C and 16/8 h light/ dark photoperiod. Pots were then transferred to the greenhouse under controlled environment and maintained until flowering and podding.

### Cloning and bioinformatics analysis of *LsAAE3*

AAE3 protein sequences from *Medicago truncatula* (GenBank: XP003599555), *Arabidopsis thaliana* (GenBank: NP190468), *Oryza sativa* (GenBank: BAS91706) and *Vigna umbellata* (GenBank: AOO87786) were retrieved from the NCBI (https://www.ncbi.nlm.nih.gov/) and subjected to multiple sequence alignment by the ClustalW function of BioEdit (Hall 1999) to identify the conserved region. Degenerate primers (Table 1) were designed from the conserved region of the AAE3 proteins to PCR amplify the *AAE3* gene from the genomic DNA of grasspea (cv. Pusa-24). The reaction mixture comprised 2.5 µL 10X buffer, 0.5 µL 10 mM dNTP, 1 µL of each primer (10 µM), 0.5 µLTaq DNA polymerase, 1 µL DNA (100 ng/µl), 19.5 µl H_2_O. PCR conditions were: initial denaturation 94°C for 5 min followed by 36 cycles of denaturation at 94°C for 1 min, annealing at 52°C for 1 min and extension at 72°C for 1 min and a final extension at 72°C for 10 min. The amplified fragment was cloned into pGEM®-T Easy vector (Promega, USA) and sequenced. The obtained sequence was trimmed to remove vector sequence and then BLAST searched to identify its similarity with the known *AAE3* genes. Two assembled but uncharacterized grasspea transcriptomes were retrieved from the GenBank (accession no. GBSS01000000, Almeida et al. 2014) and Dryad Digital Repository (http://dx.doi.org/10.5061/dryad.k9h76; Chapman 2015) and used as a subject to identify the contigs/transcript matching with the partial *LsAAE3* sequence used as a query. The identified transcript was subjected to NCBI ORF finder (https://www.ncbi.nlm.nih.gov/orffinder/) to find the Open reading frame (ORF) and translate it to protein. The predicted protein was subjected to BLAST to check its similarity with other known AAE3 proteins and ClustalW for multiple sequence alignment.

**Table 1.**
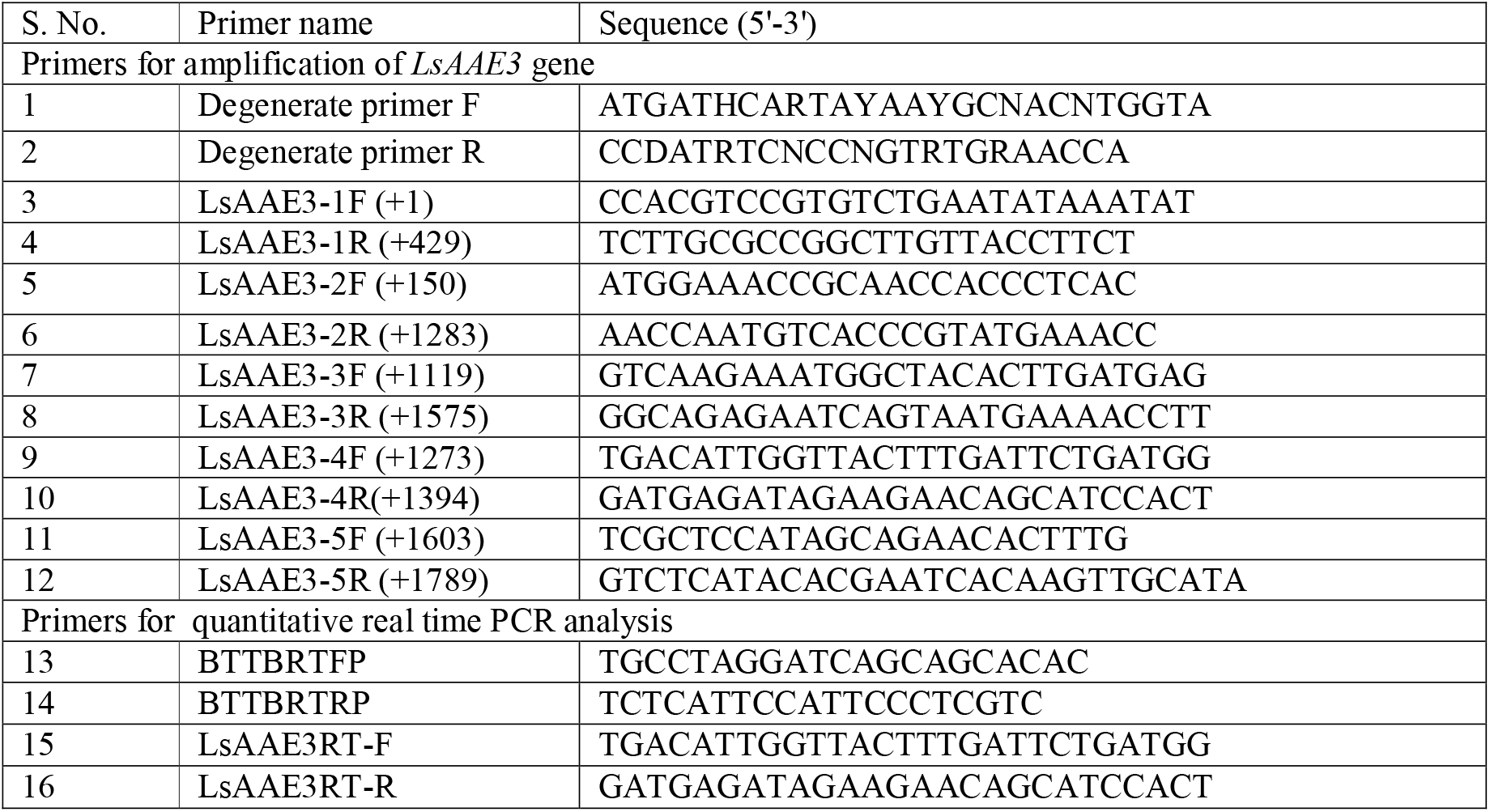
Sequences of primers used in various experiments

### Genomic PCR and determination of *LsAAE3* gene structure

Genomic DNA was isolated from young leaves of grasspea (cv. Pusa-24) using CTAB method (Doyle and Doyle 1990) with few modifications. Five sets of overlapping primers were designed from the *LsAAE3* cDNA/transcript sequence (Table 1) and used to amplify various overlapping fragments from the genomic DNA of grasspea. The PCR reaction mixture comprised 5 µL 5X buffer, 0.5 µL 10 mM dNTP, 1 µL of each primer (10 µM), 0.3 µL Phusion high-fidelity DNA polymerase 2U/µL, 1µL DNA (100 ng/µL), 16.2 µL H_2_O. PCR conditions were: initial denaturation 98°C for 30 s followed by 36 cycles of denaturation at 98°C for 10 s, annealing at 60°C for 30 s, and extension at 72°C for 1 min and a final extension at 72°C for 10 min. The amplified fragment was gel eluted using a gel extraction kit (Genetix, New Delhi) and sequenced. The obtained genomic sequences were assembled into a contiguous sequence and aligned with the *LsAAE3* cDNA sequences to identify the gene structure, number of exons and introns present in the gene sequences.

### Phylogenetic analysis and *in silico* subcellular localization study

AAE3 proteins and coding sequences (CDS) from different plant species were retrieved from the Legume Information System (LIS), RAP_DP, http://plants.ensembl.org, Phytozome, TAIR database and aligned by ClustalW in MEGA5 software (Tamura et al. 2011) with the default parameters. The phylogenetic tree was constructed using MEGA5 based on neighbor-joining method. To evaluate the reliability of the tree, a bootstrap analysis was performed with 1000 replications. The subcellular localization of *LsAAE3* was predicted through TargetP2.0 (http://www.cbs.dtu.dk/services/TargetP/), WoLFPSORT (https://wolfpsort.hgc.jp/), Mitofates (http://mitf.cbrc.jp/MitoFates/cgi-bin/top.cgi), TPpred3.0 (https://tppred3.biocomp.unibo.it/welcome/default/index), and SCLpred (https://schloro.biocomp.unibo.it/) web servers.

### Structural bioinformatics

The protein sequence of *LsAAE3* was submitted to swissmodel database (swissmodel.expasy.org/) for *in silico* structure prediction based on homologous crystal structure available in the protein data bank. The top hit template (PDB Id: 5ie2.1, oxalate-CoA ligase from *Arabidopsis thaliana*) was selected to build 3D model of LsAAE3 protein. The stereo-chemical quality of LsAAE3 protein structure was assessed by PROCHECK web server (https://servicesn.mbi.ucla.edu/PROCHECK) and the predicted structure was visualised through PyMol software.

### Estimation of seed ODAP content

Seed β-ODAP content analysed from the Mahateora and Pusa-24 variety of grasspea according to high thorough put plate-based spectrophotometer protocol (Emmrich et al. 2019). Briefly, dry seeds of Pusa-24 and Mahateora ground in a mortar and pestle and 0.5 g of fine seed powder dissolved in 10 mL of 60% ethanol (v/v) and incubated in an orbital shaker with shaking at room temperature for 16 h. The mixture was then centrifuged at 4000 rpm for 15 min and 2 mL of the supernatant was mixed with 4 mL of 3M KOH and incubated in a boiling water bath for 30 min for hydrolysis. From the hydrolysed extract, 30 µL were taken in triplicates and then mixed with 220 uL pre-diluted OPT buffer and incubated at room temperature for 30 min for colour forming reaction. The absorbance was measured at 420 nm in a microplate reader (Molecular devices Spectra Max). The standard curve is prepared using dilution series of L-DAP in the range of 0.0176 mg/mL to 0.1936 mg/mL. Nonhydrolyzed extract and blank were analysed simultaneously in the plate to reduce the background absorbance caused by other compounds in the extract.

### Total RNA isolation and real time quantitative PCR analysis

To examine the expression pattern of the *LsAAE3* gene in different tissues of grasspea, total RNA from seedling shoots (6 DAS), seedling roots (6 DAS), young leaves (10 DAS), open flowers, mature leaves (during podding stage), and early pods were isolated from the variety Pusa-24. For establishing the *LsAAE3* expression with the accumulation pattern of β-ODAP, total RNA was isolated from seedling shoots (6 DAS), seedling roots (6 DAS), young leaves (10 DAS) from the variety Pusa-24 (Medium ODAP content) and Mahateora (low ODAP content). Total RNA isolation was performed using Trizol reagent (Invitrogen, USA) (Connolly et al., 2006). DNaseI (Thermo Scientific, USA) treatment was given to remove any contaminating DNA. The quantity and purity of RNA were measured using NanoDrop 8000 Spectrophotometer (Thermo Scientific, USA). One µg of total RNA was used for reverse transcription reaction using OligodT primer and RevertAid reverse transcriptase (Thermo Scientific, USA) following the manufacturer’s protocol. qRT–PCR was performed using PowerUp SYBR Green Master Mix (Thermo Scientific, USA) in ABI 7500 real-time PCR system. PCR reaction was carried out in 10 µL using 5 µL of 2X SYBR green master mixes, 0.5 µL of each primer, 25.0 ng of cDNA from each sample. The Real-time PCR program was 50°C for 2 min, 95°C for 2 min and then 40 cycles of 95°C for 15 sec, 60°C for 1 min followed by dissociation curve program: 95°C for 15 sec, 60°C for 1 min, 95°C for 30 sec and 60°C for 15 sec to check for nonspecific amplification. All the reactions were conducted in triplicate. β*-tubulin* of *L. sativus* was used as internal control and primers for β*-tubulin* were as per Chakraborty et al. (2018). The gene-specific primers for *LsAAE3* were designed in such a way so that they anneal to exons flanking intron sequences to identify any potential genomic DNA contamination. Relative expression levels of LsAAE3 in various samples were calculated using the 2–ΔΔ*C*T method (Livak and schmittgen 2001). For establishing the LsAAE3 expression with the accumulation pattern of β-ODAP, two biological replicates were taken and each biological replicate comprised the pooled tissues from 3-5 seedlings.

### *In silico* identification of CoA dependent pathway of oxalate degradation

The cDNA sequence of *At5g14780* (Formate dehydrogenase, FDH) and *At5g17380* (Oxalyl-CoA decarboxylase, OCD) were downloaded from the TAIR database and grasspea transcriptome datasets were downloaded from the GenBank (accession no. GBSS01000000, Almeida et al. 2014) and Dryad Digital Repository (http://dx.doi.org/10.5061/dryad.k9h76; Chapman 2015) and local nucleotide database file was created using Bioedit program. The BLASTN search was performed against the local nucleotide database file and the homologs of grasspea FDH and OCD were identified with an E-value of <1e−6 and exhibiting >70% nucleotide sequence identity. Full length transcript of grasspea FDH and OCD were then subjected to NCBI ORF finder and NCBI BLASTP search to confirm its identity with the other related plant species.

## Results and discussion

### Identification of *AAE3* gene from grasspea

Due to limited availability of suitable genome sequence information and transcriptome datasets of grasspea in the public domain, we amplified the *AAE3* gene from the genomic DNA of grasspea (cv. Pusa-24) using the degenerate primers designed from the conserved region (Fig. 2a) of already characterized AAE3 proteins of other plants (Foster et al. 2012; Foster et al. 2016; Lou et al. 2016; Peng et al. 2017). A ∼443 bp fragment was PCR amplified (Fig. 2b), cloned into the TA-cloning vector (Fig. 2c) and sequenced. After trimming the vector sequences, 443 bp unique sequence having degenerate forward and reverse primers at the end of sequence was obtained (Supplementary Fig. 1). The obtained grasspea sequence was translated and then BLAST searched against the protein databases which showed 88% amino acids sequence identity with the characterised *AAE3* gene of *M. truncatula* (Foster et al. 2016), confirming that cloned sequence is a true homolog of *AAE3* gene. A BLAST search of the partial *AAE3* gene of grasspea in the two publically available, assembled but unannotated grasspea transcriptome data (Chapmen et al. 2015; Almeida et al. 2014) identified two contigs of 1849 bp (c19944_g1_i1) and 2474 bp (GBSS01000590.1) (Fig. 3a) with an E-value of 0.0, displayed >98% identity with the cloned fragment. Both the contigs shared an open reading frame (ORF) of 1566 bp, predicted to encode a protein of 521 amino acids (Fig. 3b), however, the length of 5’
s and 3’ UTR varied between the two contigs (Fig. 3a). The contig GBSS01000590.1 was found to be chimeric, in which 680 bp transcript of elsewhere locus was misassembled (Fig. 3a). Due to error in *de novo* transcriptome assembly, it is always preferable to use two and three independent transcriptome datasets and aligned them to get the consensus sequences and accurately predict the gene sequence. The deduced full length AAE3 protein of grasspea was subjected to the BLAST search to identify its similarity with the known AAE3 proteins belong to diverse organisms. The grasspea AAE3 protein displayed high sequence similarity with the AAE3 proteins of plants such as PsAAA3 of *P. sativum* (96.5%), MtAAE3 of *M. truncatula* (88%), CaAAE3 of *Cicer arietinum* (87%), and least sequence similarity with the ScAAE3 of *Saccharomyces cerevisiae* (46%) (Table 2). Therefore, we annotated the identified grasspea ORF as *LsAAE3*. Previous *in-silico* studies have shown that AAE3 protein sequences are conserved among plant species (Wang et al. 2011; Chen et al. 2017) and have been shown to function as oxalyl-CoA synthetase in *Arabidopsis, Medicago*, rice bean and rice as well as in yeast (Foster et al. 2012; Foster and Nakata 2014; Foster et al. 2016; Lou et al. 2016; Peng et al. 2017). Although the biochemical function of LsAAE3 protein has to be investigated, but its high sequence similarity with the known AAE3 proteins strongly suggest that *LsAAE3* also function as oxalyl-CoA synthetase.

**Fig. 2.**
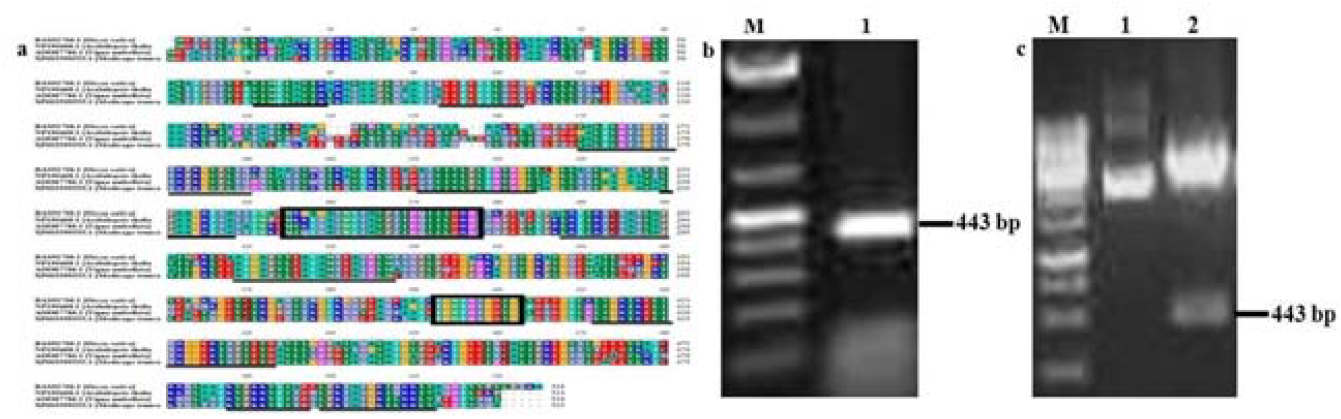
Cloning of grasspea *AAE3* homolog by PCR with degenerate primers. **a** Multiple sequence alignment of characterised *AAE3* proteins. Conserved regions were marked in black lines. Conserved regions selected for degenerate primers were marked with black boxes. **b** Agarose gel image of degenerate oligo nucleotide-primed amplicon of *AAE3* in grasspea. M: 100 bp ladder; 1: putative partial *AAE3*amplicon. **c** Restriction digestion of TA-cloning vector. M: 1 kb ladder, 1: undigested plasmid; 2: plasmid digested with *Not*I

**Fig. 3.**
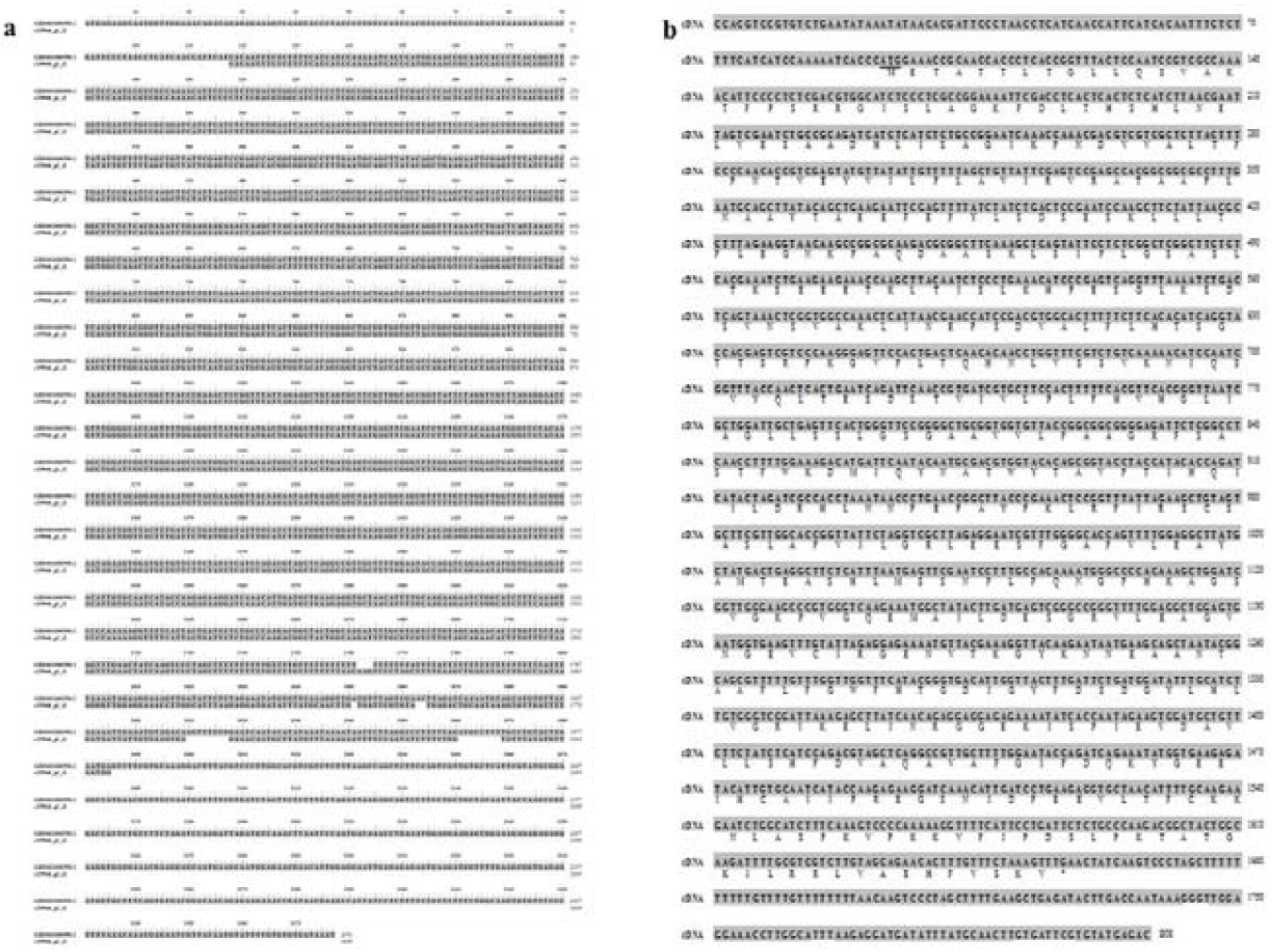
Nucleotide and deduced amino acid sequences of grasspea *LsAAE3* transcript. **a** Alignment of c19944_g1_i1 and GBSS01000590.1 contigs to determine consensus sequence of cDNA. **b** Consensus cDNA sequence and deduced amino acid sequences. The translational start codon is underlined and the translational stop codon is marked with an asterisk

**Table 2.**
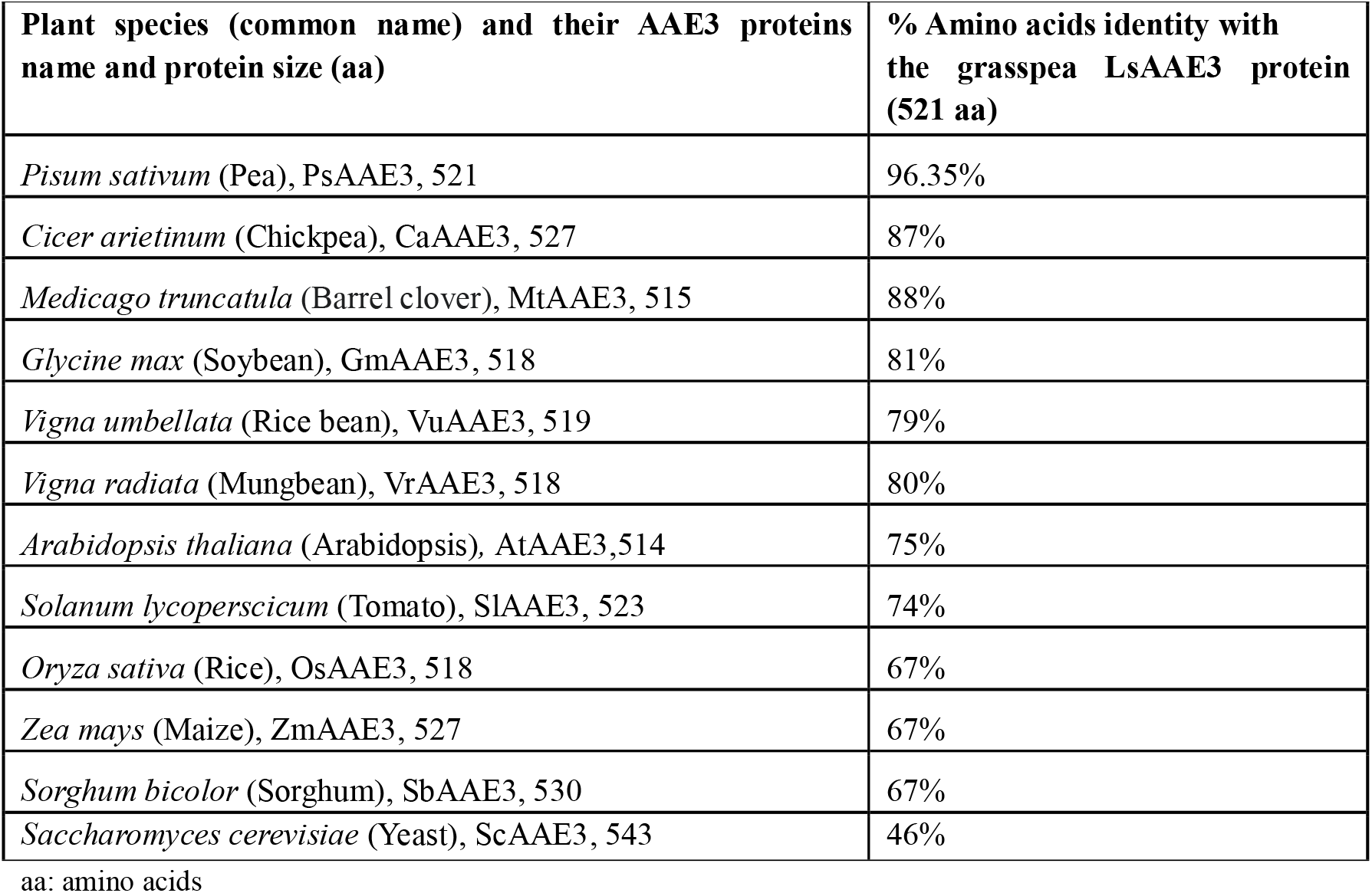
Percentage amino acid identity of grasspea LsAAE3 protein with its orthologs in plants and yeast

### Determination of *LsAAE3* gene structure

To determine the *LsAAE3* gene structure, genomic sequence of *LsAAE3* was generated using five sets of overlapping primers designed around the cDNA sequence of *LsAAE3* (Fig. 4; Table 1) and used in PCR to amplify the full length of the *LsAAE3* gene from genomic DNA of grasspea (cv. Pusa-24) and sequenced. The resultant overlapping sequences were assembled into one contiguous fragment (3,555 nucleotides). The alignment of the *LsAAE3* genomic sequence with the *LsAAE3* transcript sequence revealed the presence of four exons and three introns in the *LsAAE3* gene (Fig. 5, GenBank accession no. MH469748). The comparison of the *LsAAE3* gene structure with the predicted or characterised *AAE3* genes of dicots, monocots and yeast found that *LsAAE3* shares a highly conserved gene structure with the dicots *AAE3* but showed considerable divergence from the monocots orthologs (Table 3). Further, phylogenetic relationship analysis using protein and nucleotide sequences indicated that *AAE3* genes clustered into two groups, one consisting of *AAE3* gene of dicots and another consisting of monocots *AAE3* genes (Fig. 6a,b). The dicot *AAE3* cluster further diverged into legume and non-legume sub-clusters (Fig. 6a,b). The legume sub-cluster forms two groups with one comprising of soybean, rice bean and mungbean and the second of grasspea, pea, *Medicago* and chickpea. As expected, closely related species showed less sequence divergence than distantly related species. The close clustering of *LsAAE3* with the *Medicago AAE3* gene that has proven oxalyl-CoA synthetase activity (Foster et al. 2016) suggests that *LsAAE3* most likely functions as oxalyl-CoA synthetase.

**Fig. 4.**
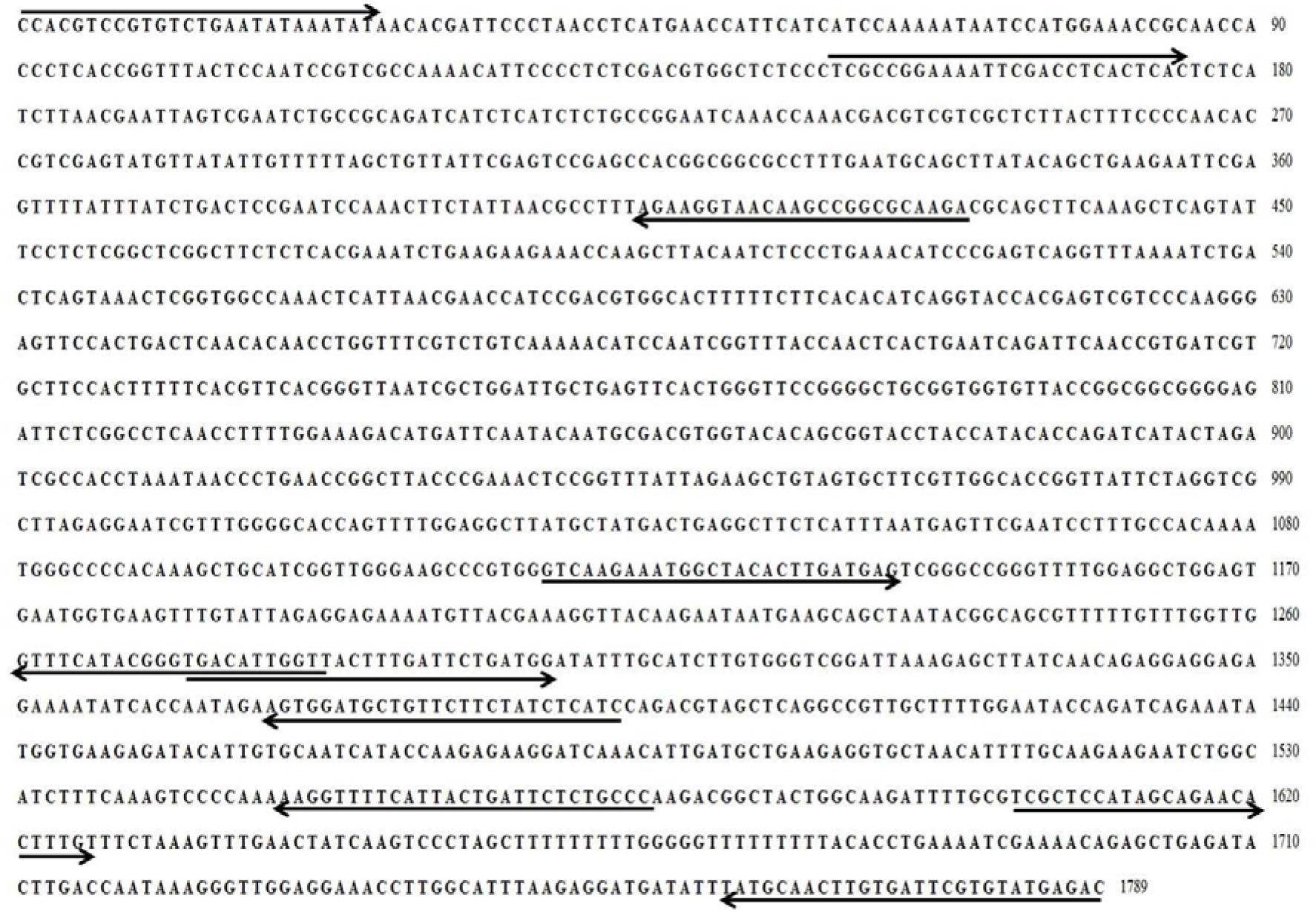
Various primers designed from the grasspea *LsAAE3* transcript. Arrows indicate the location of forward and reverse primers

**Fig. 5.**
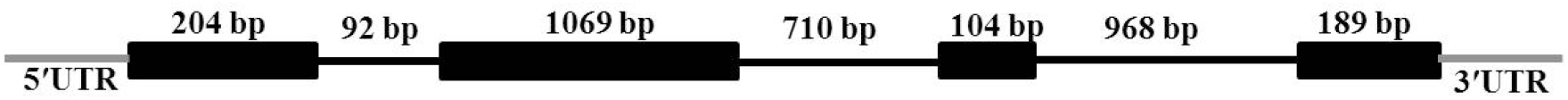
Structure of *LsAAE3* gene. The black boxes indicate exons and the black solid lines indicate introns. 5 □ and 3 □ UTR are indicated by gray lines

**Fig. 6.**
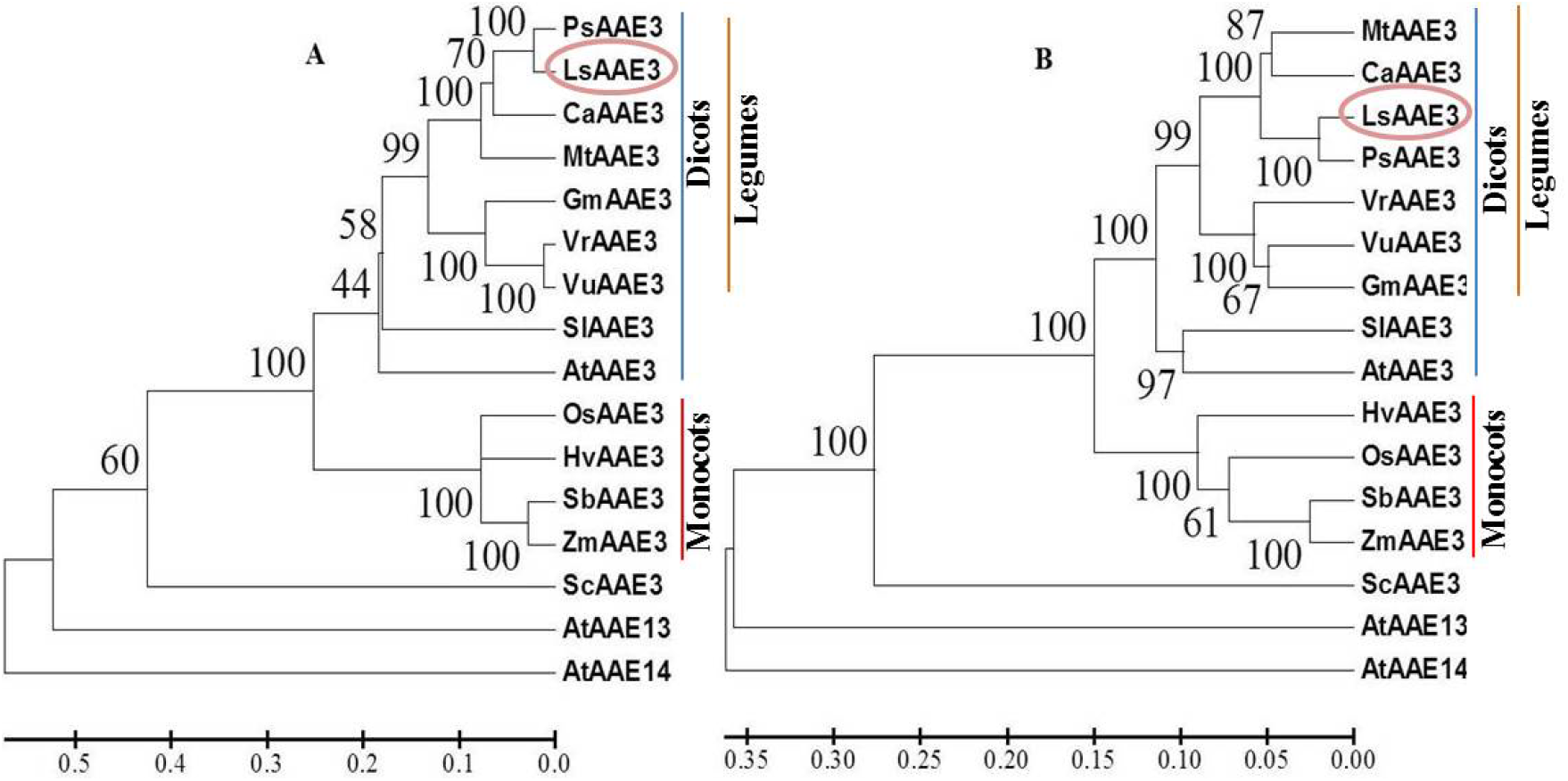
Phylogenetic relationship of *LsAAE3* and its orthologs. **a** Phylogenetic tree of *AAE3* based on nucleotide coding sequences. **b** Phylogenetic tree of *AAE3* based on proteins sequences. *AAE3* proteins were derived from *Pisum sativum* (PsAAE3), *Lathyrus sativus*(LsAAE3; GenBank: MH469748), *Cicer arietinum*(CaAAE3), *Medicago truncatula* (MtAAE3), *Glycine max* (GmAAE3), *Vigna radiata* (VrAAE3), *Vigna umbellata* (VuAAE3), *Solanum lycoperscicum* (SlAAE3), *Arabidopsis thaliana* (AtAAE3), *Oryza sativa* (OsAAE3), *Hordeum vulgare* (HvAAE3), *Sorghum bicolor* (SbAAE3), *Zea mays* (ZmAAE3), *Saccharomyces cerevisiae* (ScAAE3). AtAAE13 (TAIR:AT3G16170), AtAAE14 (TAIR:AT1G30520) of Arabidopsis were included in the tree as an outgroup

**Table 3.**
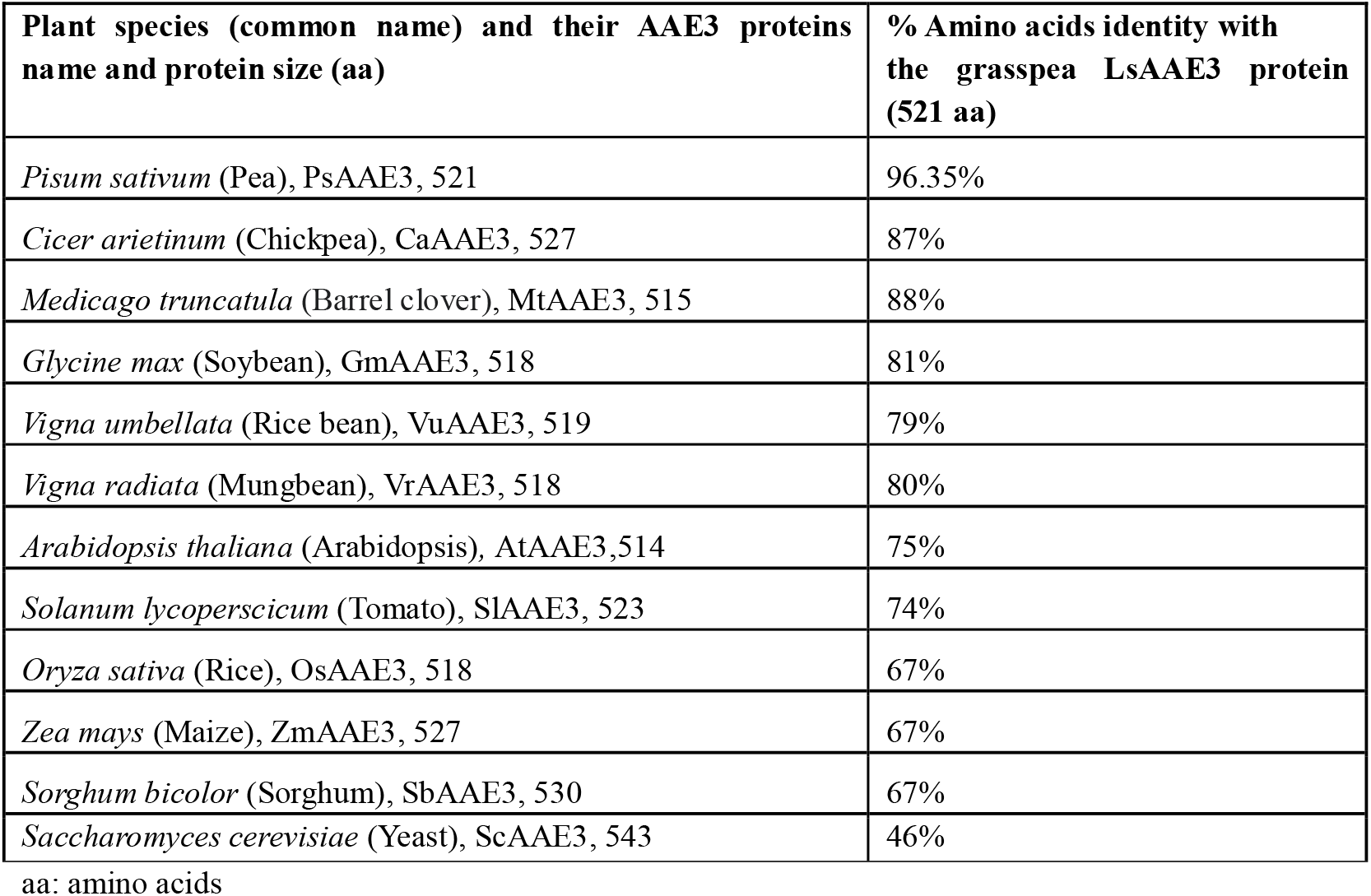
Structure of *AAE3* genes in different plant species

### Domain and sequence analysis of LsAAE3 protein

*LsAAE3* is predicted to encode a protein of 521 amino acids. Pfam search (http://pfam.xfam.org/search) using the LsAAE3 protein sequence identified two conserved domains, a large N-terminal AMP binding enzyme domain and smaller C-terminal AMP binding enzyme domain (Fig. 7). These domains are characteristics of ANL superfamily of adenylating enzymes which comprises three subfamilies: Acyl-CoA synthetases, the non-ribosomal peptide synthetases, and the firefly luciferase enzymes (Gullick 2009). In Arabidopsis, the Acyl-CoA synthetase family contains 63 different genes which occur in seven different clades (Shockey et al. 2003). *AAE3* belongs to clade VII of the family, which consists of three genes, *AAE3, AAE13*, and *AAE14* (Shockey et al. 2003). The biochemical function of all three genes in the clade VII was identified. *AAE14* gene encodes o-succinyl benzoyl-CoA ligase, *AAE13* gene encodes malonyl-CoA synthetase and *AAE3* gene encodes oxalyl-CoA synthetase (Kim et al. 2008; Chen et al. 2011; Foster et al. 2012). Since, *LsAAE3* is a grasspea ortholog of *AAE3* and shared close similarity with the AAE3 clade VII (Fig. 6; Table 2), therefore, LsAAE3 may also function as oxalyl-CoA synthetase. Further, amino acid sequence alignment of LsAAE3 with the *Arabidopsis* and *Medicago* AAE3 revealed the presence of highly conserved AMP binding domain ‘FLHTSGTTSRPK’ and Acetyl-CoA synthetase domain ‘FGWFHTGDXGXXDXXGYXXLVGRIK’ (Fig. 8) as reported in other AAE3 orthologs (Wang et al. 2011; Lou et al. 2016: Chen et al. 2017). Recently, crystal structure of *Arabidopsis AAE3* encodes an oxalyl-CoA synthetase was elucidated (Fan et al. 2016). Because of high sequence identity of LsAAE3 protein with the Arabidopsis and other *AAE3* orthologs of dicots (Fig. 6, Table 2), the three dimensional structure of *Arabidopsis AtAAE3* can serve as template for homology modelling of *LsAAE3*. Homology model structure of LsAAE3 revealed the conserved three-dimensional folding patterns similar to Arabidopsis AtAAE3 (Fig. 9a). Structure based sequence alignment revealed the residues involved in oxalate and ATP binding are highly conserved between AtAAE3 and LsAAE3 proteins (Fig. 9b). The quality of LsAAE3 protein structure was checked by Procheck web server which showed the presence of 93.7% residues in the most favoured region (red) of the Ramachandran plot (Fig. 9c), indicating predicted structure of *LsAAE3* protein is quite good (Fig. 9c). The conservation of functional key sites required for the oxalate and ATP binding as well as three-dimensional fold of *LsAAE3* as of *AtAAE3* that has shown to encode oxalyl-CoA synthetase (Foster et al. 2012), assures that *LsAAE3* encodes an oxalyl-CoA synthetase.

**Fig. 7.**
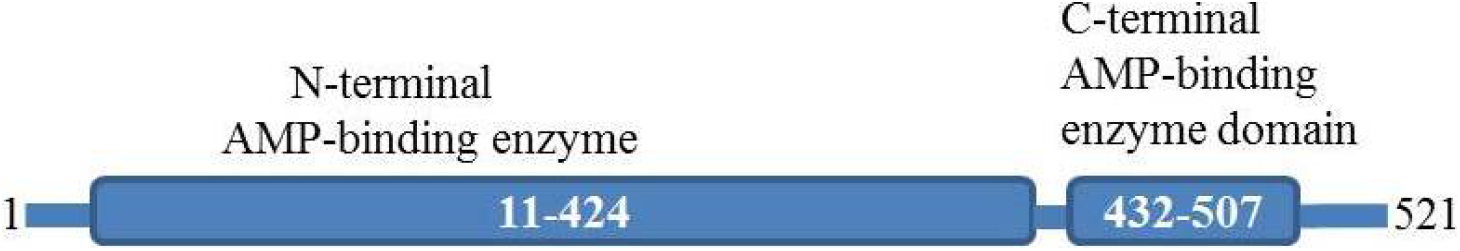
Schematic diagram of the *LsAAE3* protein showing the conserved domains.

**Fig. 8.**
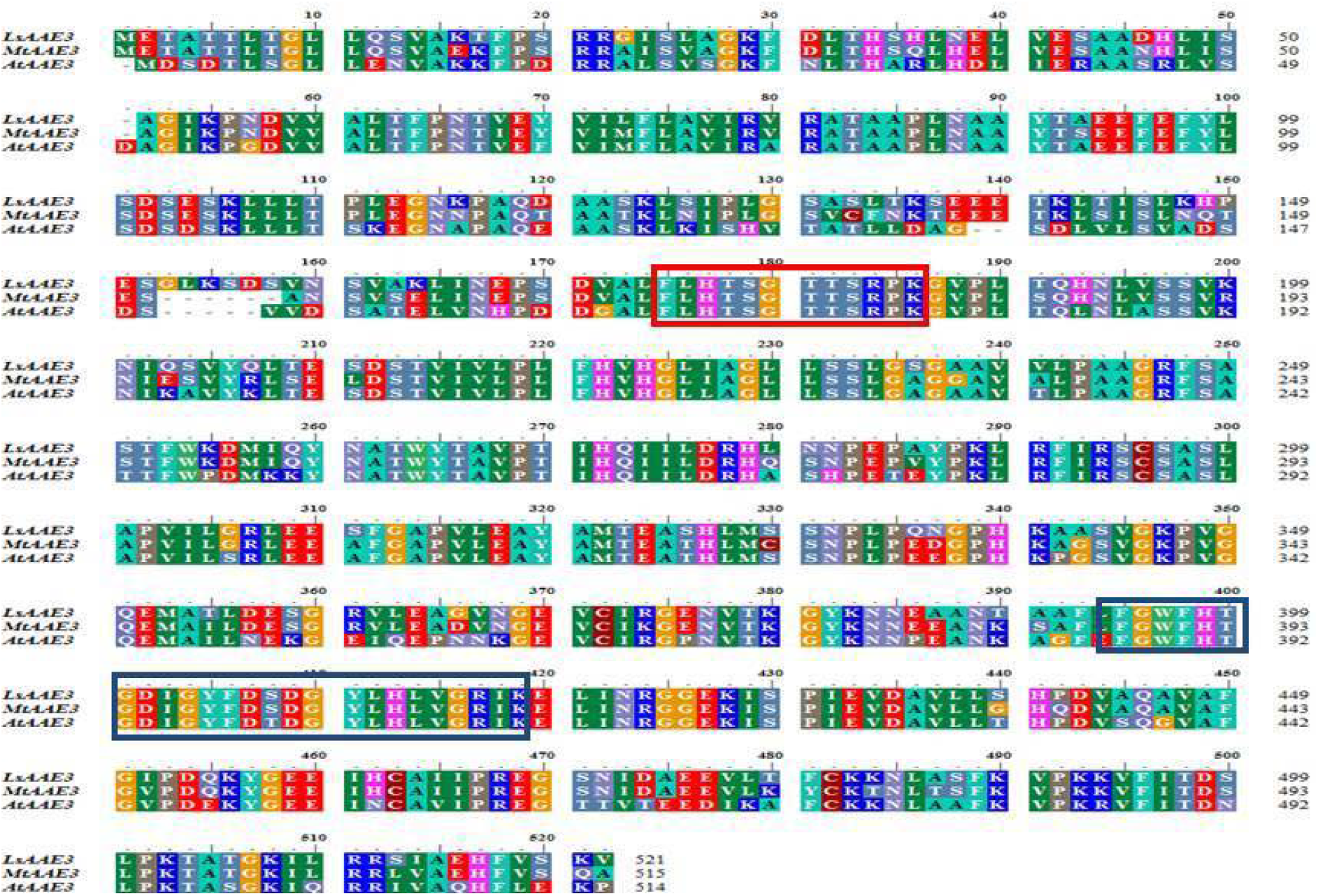
Amino acid sequence alignment of oxalyl-CoA synthetase proteins from *L. sativus* (*LsAAE3*), *A. thaliana* (*AtAAE3*), and *M. truncatula* (*MtAAE3*). The conserved AMP binding domain (red box) and Acetyl-CoA synthetase domain (blue box) are indicated

**Fig. 9.**
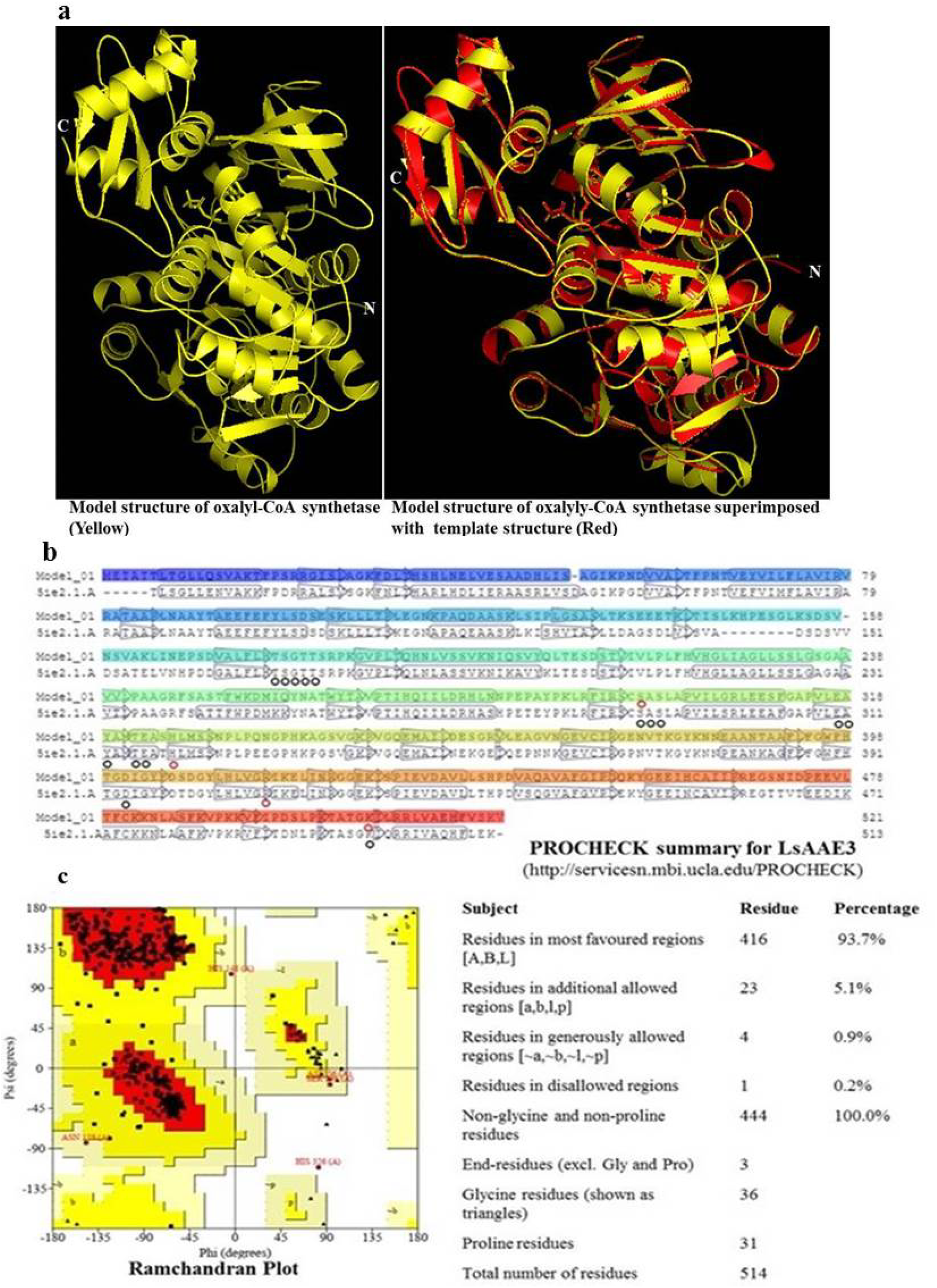
Homology-model structure of grasspea oxalyl-CoA synthetase. **a** The generated model structure of LsAAE3 protein, model structure was superimposed with the template crystal structure of oxalyl-CoA synthetase from *Arabidopsis thaliana*. **b** Structure based sequence alignment of LsAAE3 protein with the oxalyl-CoA synthetase of *A. thaliana*, the residues involved in ATP and oxalate binding are marked by black and red circles, respectively. **c** Ramachandran plot and procheck summary of LsAAE3 model structure.

### *In-silico* subcellular localization of LsAAE3 protein

The subcellular localization study of LsAAE3 revealed its non-organellar localization as well localization in multiple compartments with a very low-reliability score. However, none of the results from different web servers study have predicted the localization of LsAAE3 in chloroplast or mitochondria (Table 4). The results are in agreement with the previous work reported in Medicago, Arabidopsis, maize, rice and rice bean where localization of AAE3 proteins were reported to be in the cytoplasm based on the transient or stable expression of GFP-tagged *AAE3* proteins in the plant cell (Foster et al. 2012; Foster et al. 2016; Lou et al. 2016; Peng et al. 2017). Previous studies have suggested the biosynthesis of β-ODAP in the chloroplast or mitochondria (Ikegami et al. 1992; Ikegami et al. 1993; Xu et al. 2017). However, we did not find any chloroplast or mitochondrial targeting peptide signal in the LsAAE3 protein (Table 4), suggesting *LsAAE3* encoded oxalyl-CoA synthetase might not be involved in β-ODAP biosynthesis of grasspea.

**Table 4.**
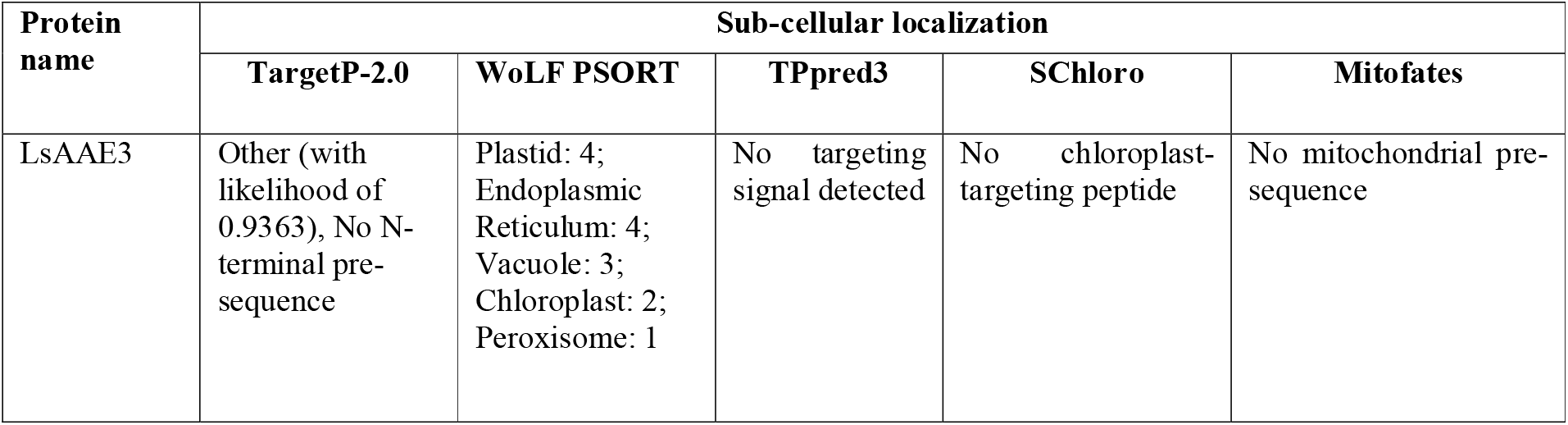
Summary of possible sub-cellular localization of LsAAE3 protein of grass pea

### Expression analysis of *LsAAE3*

To estimate the relative expression level of *LsAAE3* gene in different tissues such as seedling shoot and root, leaf at seedling and podding stage, flower, and green pod (containing immature seeds), qRT–PCR assay was performed. The qRT–PCR detected transcripts of *LsAAE3* gene in different tissues at varying level with expression being highest in the leaf and lowest in the root at the seedling stage (Fig. 10a). Based on report from *Arabidopsis* and *Medicago* eFP browser (http://bar.utoronto.ca/efp2/Arabidopsis/Arabidopsis_eFPBrowser2.html), (http://bar.utoronto.ca/efpmedicago/cgi-bin/efpWeb.cgi), *AAE3* gene of *Medicago* and *Arabidopsis* is also expressed in broad range of tissues. Similarly, *AAE3* orthologue in maize also showed wider expression patterns in tissues like root, stem, leaf, husk, silk, tassel, ear, and kernel (Wang et al. 2011). Further, to establish the possible link of *AAE3* encoded oxalyl-CoA synthetase with the β-ODAP biosynthesis, we compared the expression level of *LsAAE3* in medium (cv. Pusa-24 (ODAP content 0.2 to 0.4%) and low (cv. Mahateora) (ODAP content 0.04 to 0.05%) ODAP containing varieties of grasspea. Seedling stage, especially upto 10 days after sowing has been reported to accumulate highest amount of β-ODAP in comparison to other developmental stages (Zhang et al. 2005; Jiao et al. 2006). Therefore, six days old shoot and root and 10 days old young leaf of Pusa-24 and Mahateora were taken for relative expression analysis. Results of qRT–PCR are presented in Fig. 10b, taking *LsAAE3* transcript level of Pusa-24 as a calibrator. The *LsAAE3* transcript levels in the root were comparable between Pusa-24 and Mahateora. However, *LsAAE3* transcript levels in the young shoot and leaf were 5.1 and 3.71 folds higher in Mahateora than Pusa-24, respectively (Fig. 10b). Previous study reported that young seedlings, especially six days old young shoots, accumulated highest content of β-ODAP (Jiao et al. 2006). Similar to the accumulation pattern of β-ODAP, the maximum difference in expression of *LsAAE3* between two varieties was observed in the six days old young shoots. Since *LsAAE3* is predicted to encode an oxalyl-CoA synthetase, reported to be involved in biosynthesis of β-ODAP (Malathi et al., 1970), the medium-ODAP containing varieties are expected to have higher *LsAAE3* expression in comparison to low-ODAP containing varieties. In contrast, we obtained higher expression of *LsAAE3* in the low-ODAP than medium-ODAP containing variety suggesting *LsAAE3* encoded oxalyl-CoA synthetase might not be directly involved in biosynthesis of β-ODAP, instead, it appears to negatively regulate the β-ODAP biosynthesis pathway. This observation is supported by a series of evidence. Firstly, oxalic acid is the precursor of β-ODAP (Kumar et al., 2016). As per Srivastava et al. (2015), the oxalate content in Pusa-24 is 1.44 mg/g and Mahateora is 1.42 mg/g, which is almost similar. However, these varieties differ in terms of ODAP content (0.04 to 0.05% in Mahateora and 0.2 to 0.4% in Pusa-24), it means lower ODAP containing variety utilises less oxalate for the synthesis of β-ODAP compared to high ODAP containing variety. Secondly, excessive accumulation of oxalate has deleterious effects on growth and development of the plants (Foster et al. 2016; Yang et al. 2018), therefore, similar to other plant species, grasspea might also have evolved a CoA-dependent pathway of oxalate degradation and possibly utilises this pathway when the demand of oxalate for the β-ODAP synthesis is less. Thirdly, all the characterised AAE3 orthologs have been shown to be involved in CoA-dependent pathway of oxalate degradation (Foster et al. 2012; Foster et al. 2016; Lou et al. 2016; Peng et al. 2017). Fourthly, previous studies have suggested the biosynthesis of β-ODAP in the chloroplast or mitochondria (Ikegami et al. 1992; Ikegami et al. 1993; Xu et al. 2017). However, our *in silico* localization study did not find any chloroplast or mitochondrial targeting peptide signal in the LsAAE3 protein (Table 4). Finally, previous studies have shown the oxalate induced up-regulation of AAE3 orthologs (Foster et al. 2016; Lou et al. 2016). Based on these evidences and our expression study, we hypothesized that reduced synthesis of β-ODAP induces LsAAE3 expression to metabolised unutilised oxalate via CoA-dependent pathway of oxalate degradation.

**Fig. 10.**
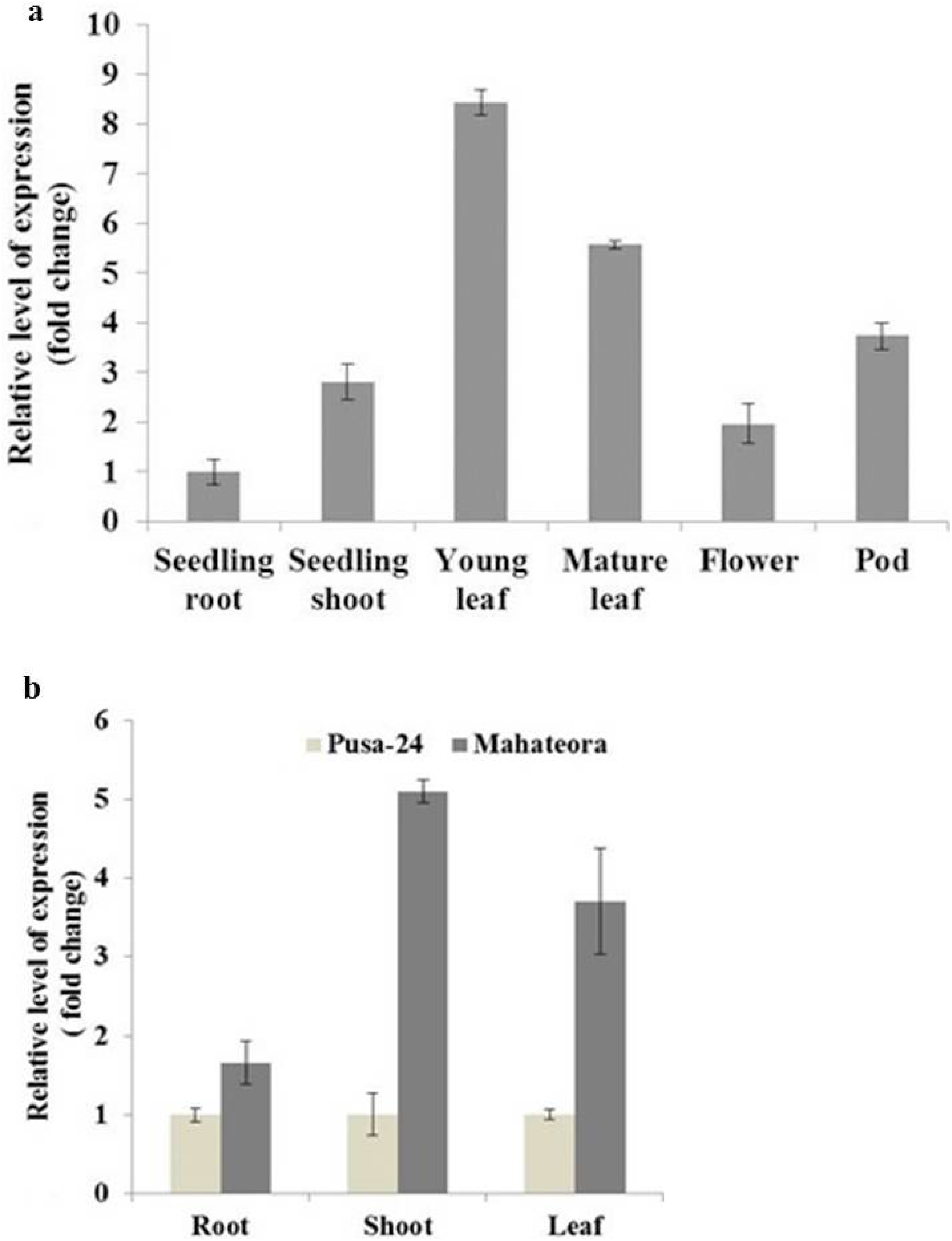
Analysis of *LsAAE3* gene transcript. **a** Bar chart showing relative expression of *LsAAE3* in tissues like seedling root, seedling shoot, seedling leaf, mature leaf, flower and green pod (cv. Pusa-24). **b** Bar chart showing relative expression of *LsAAE3* gene transcript in seedling tissues like six days old root, shoot, ten days old young leaf of Pusa-24 and Mahateora. Error bar represents Standard Error of six technical replicates and two biological replicates. Pusa-24 was taken as calibrator

### Identification of CoA-dependent pathway of oxalate degradation in grasspea

To support that *LsAAE3* is indeed an oxalyl-CoA synthetase specifically involved in CoA-dependent pathway of oxalate degradation, it is necessary to explore the other components of the pathway to unequivocally establish the existence of such pathway in grasspea. We searched the other genes encoding enzymes of CoA-dependent pathway of oxalate degradation, namely, Oxalyl-CoA decarboxylase (*OCD*) and Formate dehydrogenase (*FDH*) homolog in publically available transcriptome database of grasspea using known *FDH, At5g14780* and *OCD, At5g17380* gene of *Arabidopsis* as a query (Foster et al. 2012). For the grasspea *FDH* homolog, two transcripts of 1619 bp (c29619_g1_i3; Chapmen 2015) and 2059 bp (GBSS01000739.1; Almeida et al. 2014) were identified that shared 75% nucleotide sequence identity with the *Arabidopsis AtFDH* gene. Both the grasspea transcripts predicted to have an ORF of 1137 bp, encode a protein of 378 amino acids. A BLASTP search using grasspea *FDH* as a query identified its homologs in *Arabidopsis, Medicago* and rice bean, shares 81%, 89% and 81% sequence identity with the *AtFDH, MtFDH* and *VuFDH* proteins, respectively. Similarly, for the *OCD* gene, two transcripts of 2984 bp (c27876_g1_i2; Chapmen 2015) and 2412 bp (GBSS01001545.1; Almeida et al. 2014) were identified displayed 73 to 74% nucleotide sequence identity with the *AtOCD* gene of *Arabidopsis*. Both these transcripts are predicted to have an ORF of 1716 bp, encode a protein of 571 amino acids. BLASTP search using the putative *OCD* protein of grasspea identified its homologs in *A. thaliana*, maize, *M. truncatula*, shared 78%, 74% and 87% amino acid identity with the *AtOCD, ZmOCD, MtOCD* proteins, respectively. The presence of transcript of Oxalyl-CoA decarboxylase and Formate dehydrogenase gene in the two independent transcriptome database of grasspea as well as high similarity of deduced protein with the known genes indicate the presence of functionally active CoA-dependent pathway of oxalate degradation in grasspea. Similar approach has also been used to support the existence of CoA-dependent pathway of oxalate degradation in *Arabidopsis* and *Medicago* (Foster et al. 2012; Foster et al. 2016).

### Conclusions

In this study, we report cloning and characterization of *AAE3* homolog from grasspea. encoding an oxalyl-CoA synthetase. We also found the transcriptionally active components of CoA-dependent pathway of oxalate degradation in grasspea. Based on expression analysis and available literature, we hypothesized that *LsAAE3* encoded oxalyl-CoA synthetase possibly negatively regulates β-ODAP biosynthesis *via* CoA-dependent pathway of oxalate degradation. Further study is required to create knockout of *LsAAE3* gene to understand how the two oxalate metabolic pathways i.e. β-ODAP biosynthesis and CoA-dependent pathway of oxalate degradation is regulated in grasspea.

## Acknowledgments

The authors are thankful to the Indian Council of Agricultural Research (ICAR) for providing financial support.

## Author contributions

Neetu Singh Kushwah conceived and designed the experiments, taking valuable suggestions and scientific inputs from Shanmugavadivel, P.S. and Alok Das. Neetu Singh Kushwah performed the experiments. Data were analysed by Neetu Singh Kushwah and Shanmugavadivel, P.S. The manuscript was prepared by Neetu Singh Kushwah in consultation with Narendra Pratap Singh, Archana Singh, Meenal Rathore, Alok Das Shanmugavadivel, P.S. All authors read and approved the final version of the manuscript.

## Conflicts of Interest

The authors declare no competing financial/personal interest related to this study.

## Abbreviations

AAE3: Acyl activating enzyme3
β-ODAP: β-*N*-oxalyl-L-α,β-diaminopropionic acid
BLAST: Basic Local Alignment Search Tool
FDH: Formate dehydrogenase
OCD: Oxalyl-CoA decarboxylase
qRT-PCR: Quantitative real-time PCR
L-DAP: L-2,3-diaminopropionic acid

## Notes

### Competing Interest Statement

The authors have declared no competing interest.

## References

Abd El Moneim AM, Van Dorrestein B, Baum M, Ryan J, Bejiga G (2001) Role of ICARDA in improving the nutritional quality and yield potential of grasspea (Lathyrus sativus L.), for subsistence farmers in dry areas. Lathyrus Lathyrism Newsletter 2(2):55–58.

Adsule RN, Barat GK (1977) Occurrence of oxalyl-CoA synthetase in Indian pulses. Experientia 33(4):416–417

Almeida NF, Leitão ST, Krezdorn N, Rotter B, Winter P, Rubiales D, and Patto, MCV (2014) Allelic diversity in the transcriptomes of contrasting rust-infected genotypes of Lathyrus sativus, a lasting resource for smart breeding. BMC Plant Biol 14(1):1–15

Campbell CG (1997) Grass pea. Lathyrus sativus L. Promoting the conservation and use of underutilized and neglected crops. Rome: Institute of Plant Genetics and Crop Plant Research, Gatersleben/International Plant Genetic Resources Institute. https://www.bioversityinternational.org/e-library/publications/detail/grass-pea-lathyrus-sativus-l/

Campbell CG, Briggs, CJ (1987) Registration of low neurotoxin content Lathyrus germplasm LS 8246. Crop Sci 27(4):821–821

Chakraborty S, Mitra J, Samanta MK, Sikdar N, Bhattacharyya J, Manna A et al, (2018) Tissue specific expression and in-silico characterization of a putative cysteine synthase gene from Lathyrus sativus L. Gene Expr Patterns 27:128–134

Chapman MA (2015) Data from: Transcriptome sequencing and marker development for four underutilized legumes. Dryad, Dataset, https://doi.org/10.5061/dryad.k9h76

Chen H, Kim HU, Weng H (2011) Malonyl-CoA synthetase, encoded by ACYL ACTIVATING ENZYME13, is essential for growth and development of Arabidopsis. The Plant Cell, 23(6):2247–2262

Chen WW, Fan W, Lou HQ, Yang JL, Zheng SJ (2017) Regulating cytoplasmic oxalate homeostasis by Acyl activating enzyme3 is critical for plant Al tolerance. Plant Signal Behav 12(1):e1276688

Cheng N, Foster J, Mysore KS, Wen J, Rao X, Nakata PA (2018) Effect of Acyl Activating Enzyme (AAE) 3 on the growth and development of Medicago truncatula. Biochem Bioph Res Co. 505(1):255–260

Connolly MA, Clausen PA, Lazar JG (2006) Preparation of RNA from plant tissue using trizol. Cold Spring Harbor Protocols, 2006 (1), pdb–prot4105

Doyle JJ, Doyle JL (1990) Isolation of plant DNA from fresh tissue. Focus, 12(13):39–40

Emmrich PMF., Rejzek M., Hill L., Brett P., Edwards A., Sarkar A., Field R.A., Martin C., Wang T.L. (2019) Linking a rapid throughput plate-assay with high-sensitivity stable-isotope label LCMS quantification permits the identification and characterisation of low β-L-ODAP grass pea lines. BMC Plant Biology, 19:489.

Fan M, Xiao Y, Li M, Chang W (2016) Crystal structures of Arabidopsis thaliana oxalyl-CoA synthetase essential for oxalate degradation. Mol. Plant 9(9):1349–1352

Fikre A, Korbu L, Kuo YH, Lambein F (2008) The contents of the neuro-excitatory amino acid β-ODAP (β-N-oxalyl-l-α, β-diaminopropionic acid), and other free and protein amino acids in the seeds of different genotypes of grass pea (Lathyrus sativus L.). Food Chem. 110(2):422–427

Foster J, Nakata PA (2014) An oxalyl-CoA synthetase is important for oxalate metabolism in Saccharomyces cerevisiae. FEBS letters, 588(1):160–166

Foster J, Kim HU, Nakata PA (2012) A previously unknown oxalyl-CoA synthetase is important for oxalate catabolism in Arabidopsis. The Plant Cell 24(3):1217–1229

Foster J, Luo B, Nakata PA (2016) An oxalyl-CoA dependent pathway of oxalate catabolism plays a role in regulating calcium oxalate crystal accumulation and defending against oxalate-secreting phytopathogens in Medicago truncatula. PLoS One 11(2), e0149850.

Giovanelli J. and Tobin N.F. (1961) Adenosine triphosphate-and coenzyme A-dependent decarboxylation of oxalate by extracts of peas. Nature 190: 1006–1007.

Gulick AM (2009) Conformational dynamics in the Acyl-CoA synthetases, adenylation domains of non-ribosomal peptide synthetases, and firefly luciferase. ACS Chem Biol 4(10):811–827.

Hall TA (1999) BioEdit: a user-friendly biological sequence alignment editor and analysis program for Windows 95/98/NT. Nucleic acids symposium series 41:95–98

Ikegami F, Horiuchi S, Kobori M, Morishige I, Murakoshi I (1992) Biosynthesis of neuroactive amino acids by cysteine synthases in Lathyrus latifolius. Phytochemistry 31(6):1991–1996

Ikegami F, Ongena G, Sakai R, Itagaki S, Kobori M, Ishikawa T et al (1993) Biosynthesis of β-(isoxazolin-5-on-2-yl)-alanine, the precursor of the neurotoxin β-N-oxalyl-L-α, βdiaminopropionic acid, by cysteine synthase in Lathyrus sativus. Phytochemistry 33:93–98

Jiao CJ, Wang CY, Li FM, Li ZX, Wang YF (2006) Accumulation pattern of toxin beta-ODAP during lifespan and effect of nutrient elements on beta-ODAP content in Lathyrus sativus seedlings. J Agric Sci 144:369–375

Kim HU, Oostende CV, Basset GJ, Browse J (2008) The AAE14 gene encodes the Arabidopsis o-succinylbenzoyl-CoA ligase that is essential for phylloquinone synthesis and photosystem-I function. Plant J 54(2):272–283

Kumar S, Bejiga G, Ahmed, S, Nakkoul H, Sarker A (2011) Genetic improvement of grass pea for low neurotoxin (β-ODAP) content. Food Chem Toxicol 49(3):589–600

Kumar V, Chattopadhyay A, Ghosh S, Irfan M, Chakraborty N, Chakraborty S, Datta A (2016) Improving nutritional quality and fungal tolerance in soya bean and grass pea by expressing an oxalate decarboxylase. Plant Biotechnol J, 14:1394–1405.

Lambein F, Travella S, Kuo YH, Van Montagu M, and Heijde M (2019) Grass pea (Lathyrus sativus L.): orphan crop, nutraceutical or just plain food? Planta 250:821–838

Livak K.J, Schmittgen TD (2001) Analysis of relative gene expression data using real-time quantitative PCR and the 2−ΔΔCT method. Methods 25(4):402–408

Lou HQ, Fan W, Xu JM, Gong YL, Jin JF, Chen WW et al (2016) An oxalyl-CoA synthetase is involved in oxalate degradation and aluminum tolerance. Plant Physiol, 172(3):1679–1690

Malathi K, Padmanaban G, Sarma PS (1970) Biosynthesis of β-N-oxalyl-l-α, β-diaminopropionic acid, the Lathyrus sativus neurotoxin. Phytochemistry 9(7):1603–1610

Peng C, Liang X, Liu EE, Zhang JJ, Peng XX (2017) The oxalyl-CoA synthetase-regulated oxalate and its distinct effects on resistance to bacterial blight and aluminium toxicity in rice. Plant Biol 19(3):345–353

Shockey JM, Fulda, MS (2003) Arabidopsis contains a large superfamily of acyl-activating enzymes. Phylogenetic and biochemical analysis reveals a new class of acyl-coenzyme A synthetases. Plant Physiol 132(2):1065–1076

Srivastava RP, Singh J, Singh NP, Singh D (2015). Neurotoxin and other anti-nutrients of khesari (Lathyrus sativus) genotypes and their reduction by water soaking and dehusking. Ind J Agric Biochem 28(2):172–177

Tamura K, Peterson D, Peterson N, Stecher G, Nei M, Kumar S (2011) MEGA5: molecular evolutionary genetics analysis using maximum likelihood, evolutionary distance, and maximum parsimony methods. Mol Biol Evol 28(10):2731–2739

Wang G, Sun X, Wang G, Wang F, Gao Q, Sun X et al (2011) Opaque7 encodes an acyl-activating enzyme-like protein that affects storage protein synthesis in maize endosperm. Genetics 189(4):1281–1295

Xu Q, Liu F, Chen P, Jez JM, Krishnan HB (2017) β-N-Oxalyl-L-α, β-diaminopropionic acid (β-ODAP) content in Lathyrus sativus: the integration of nitrogen and sulfur metabolism through β-cyanoalanine synthase. Int J Mol Sci 18(3):526.

Yang J, Fu M, Ji C, Huang Y, Wu Y (2018) Maize Oxalyl-CoA Decarboxylase1 degrades oxalate and affects the seed metabolome and nutritional quality. The Plant Cell 30(10):2447–2462

Zhang DW, Xing GM, Xu H, Yan ZY, Wang CY, Wang YF, Li ZX (2005) Relationship between oxalic acid and the metabolism of beta-N-oxalyl-alpha, beta-diaminopropionic acid (ODAP) in grass pea (Lathyrus sativus L.). Isr J Plant Sci 53:89–96

