## Supplementry Fig 1 for "Identification of oxalyl-CoA synthetase gene (*LsAAE3*) and its regulatory role in β-ODAP biosynthesis in grasspea (*Lathyrus sativus* L.)"

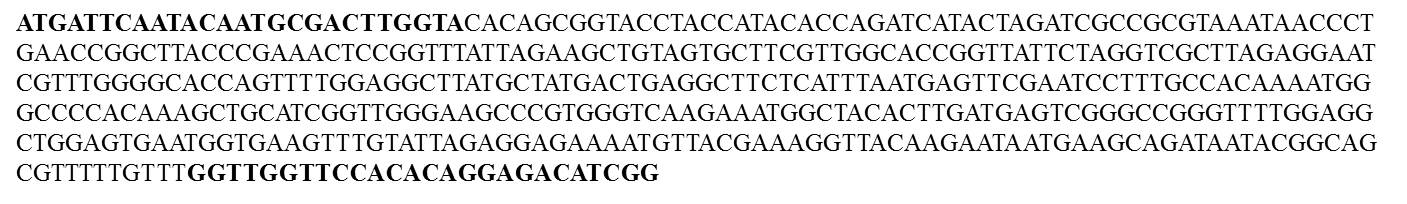

**Supplementary Fig. 1** Sequence of degenerate oligonucleotide-primed PCR amplified product. Primers sequences of forward and reverse degenerate primers are marked with bold.
